# CanID-PCR: A quick and low-cost PCR tool to identify *Candida* species on gDNA directly extracted from positive blood bottles

**DOI:** 10.1101/2025.05.13.653919

**Authors:** Hassan Badrane, Cornelius J. Clancy, M. Hong Nguyen

## Abstract

*Candida* bloodstream infection carries a high mortality. *Candida* species prone to antifungal resistance (e.g. *C. auris* and *C. glabrata*) are rising, and delay in species identification might adversely affect patients’ outcomes. Current workflow by clinical microbiology laboratory requires ∼24 hours for Candida speciation, time to allow for growth of isolates on the agar plates for testing. We established a simple and inexpensive PCR tool, named CanID-PCR, to speciate *Candida* directly from positive blood bottles. We selected *Candid*a *ACT1* gene, which has an intron with varying length that enables the identification of 10 common *Candida* species. The tool was optimized for gDNA extracted directly from positive blood culture bottles. We showed, by testing positive blood cultures from 64 unique patients, that our tool detects the same species as the microbiology lab in 97% (62/64) of samples. The 2 samples with mismatched results were due to high similarity between *Candida metapsilosis* and *C. parapsilosis*, and failure of CanID-PCR to identify *C. tropicalis* in a patient with *C. albicans/C. tropicalis* fungemia. On the other hand, CanID-PCR identified a second species (*C. fabianii*) in a patient with *C. parapsilosis* fungemia that was missed by the microbiology laboratory. In conclusion, our inexpensive tool was accurate for rapid speciation of *Candida* directly from blood culture bottles, which could be valuable for clinical and/or research laboratories.

## INTRODUCTION

*Candida* is the most common cause of nosocomial fungal bloodstream infection (BSI), which carries a high fatality rate. The most common causes of Candida BSI are *Candida albicans, C. glabrata, C. parapsilosis, C. tropicalis* and *C. krusei*, (1) although other *Candida* species such as *C. auris* are emerging (2). Rapid and accurate *Candida* species identification is important in clinics since it informs the likely resistance profile and provides timely antifungal therapy and (3, 4). It is also important for epidemiological and bench research purposes.

Several techniques, including phenotypic, proteomic and molecular methods, have been developed to speciate *Candida* isolates in clinical laboratories (5). Conventional phenotypic/biochemical methods are laborious, lengthy and suffer from lack of accuracy. Molecular techniques have been applied to a limited number of *Candida* species and/or relied on the use of multiple targets, which made them impractical and complex. Finally, commercial instruments (like the widely used matrix-assisted laser desorption ionization– time-of-flight mass spectrometry; MALDI-TOF MS) mostly provide rapid and accurate results, but require a high upfront investment (6). All above methods rely on recovery of the fungal isolate from sample culture, which usually takes an additional 24 hours. Recently, molecular techniques such as multiplex PCR of positive blood culture samples have been introduced to the clinics (e.g. Biofire or cobas eplex). However, these systems require costly instrument and reagent purchases, and integration with existing lab systems, which can be complicated. From a practical point of view, these systems are valuable to bacterial bloodstream infection (BSI), but less so for fungal BSI because the prevalence of *Candida* and other fungi is much less common than bacteria.

Here we describe a simple, affordable and quick PCR tool to identify *Candida* species from genomic DNA (gDNA) directly extracted from positive blood bottles, without the need for the extra culture on a petri dish to isolate the sample. This tool yields results in ∼ 8 hours, using regular basic laboratory techniques, which makes it convenient for research and even some clinical laboratories.

## MATERIALS AND METHODS

### Strains & Growth Conditions

The sources of *C. albicans*, *C. glabrata*, *C. tropicalis*, *C. krusei*, *C. tropicalis*, *C. parapsilosis*, *C. lusitaniae*, *C. auris*, and *C. dubliniensis* are listed in TABLE 1. *C. fabianii* was recovered from a patient’s blood culture in our researchlaboratory where the speciation was confirmed using MALDI-TOF MS. When needed, strains were grown in Yeast-Peptone-Dextrose (YPD), Synthetic Defined (SD), or CHROMAgar Candida (CHRCA) media at 30°C or 37°C.

**TABLE 1.** Candida reference isolates used in this study.

| **Species** | **Strain** | **Reference** | **ACT1 Band size**<br>**(bp)** |
| --- | --- | --- | --- |
| *C. orthopsilosis* | CDC317 | (18) | 519 |
| *C. auris* | B8441 (AR-0387) | (19) | 589 |
| *C. lusitaniae* | ATCC 66035 | (20) | 644 |
| *C. parapsilosis* | ATCC 22019 | (21) | 733 |
| *C. dubliniensis* | ATCC MYA-646 | (22) | 767 |
| *C. albicans* | SC5314 | (23) | 793 |
| *C. tropicalis* | ATCC 20336 | (24) | 824 |
| *C. fabianii* | 62862-CF60A | Badrane et al. (PC) | 861 |
| *C. glabrata* | CBS138 | (25) | 1,012 |
| *C. krusei* | ATCC 6258 | (26) | 1,144 |

### gDNA extraction directly from positive blood bottles

Two to 10 ml of broth from positive blood culture bottles were treated twice with 2x volume of Alkali Solution (0.1M Sodium Citrate, 1M Sodium Hydroxide) (7) for 10 min with rotation at medium speed. The sample was then centrifuged at 3155 x*g* for 5 min, and the pellet resuspended in 1ml of sterile nuclease-free water in an Eppendorf tube. After centrifugation, the pellet was resuspended in 600 µl extraction buffer (CLS-Y) from FastDNA Kit (MPBIO, San Diego, CA, U.S.A.). Cell disruption in a beadbeater (Biospec Products, Bartlesville OK) and DNA extraction were performed per recommendations of the respective manufacturer. DNA was eluted from the binding matrix with 200 µl of DES buffer. The eluted DNA was mixed with equal volume of membrane binding solution from the Wizard SV Gel and PCR clean-up System (Promega, Madison, WI, U.S.A.). Finally, gDNA was eluted in 40 µl nuclease-free water.

### Fingerprinting using RAPD

We first designed 10-mer RAPD primers for DNA fingerprinting of *Candida* isolates (TABLE 2). RAPD was then used on gDNA extracted using standard methods from *Candida* clones picked from blood-bottles sample spread on SD Agar. RAPD PCR was performed with the selected primers using 5PRIME Hotmaster Taq DNA polymerase (QuantaBio, Beverly, MA, U.S.A.), and set up in a 25 µl reaction volume as follows: 0.5-1 µl gDNA, 2 µl of 5µM RAPD primer, 0.5 µl of 10 mM dNTP mix, 2.5 µl of 10x Taq buffer, 0.5 µl Enzyme (5U/µl), and 25 µl nuclease-free molecular biology grade water qsp. The PCR was run on a MJ Research Diad thermocycler (Biorad, Hercules, CA, U.S.A.) using the following program: (i) an initial denaturation at 94°C for 1 min; (ii) 35 cycles of : 94°C for 15 sec, 32-35°C (depending on the primer) for 30 sec, 68°C for 40 sec; and (iii) a final extension at 68°C for 10 min. PCR products were run on a 1% agarose gel in 1x TAE for 1.5 hours.

**TABLE 2.**
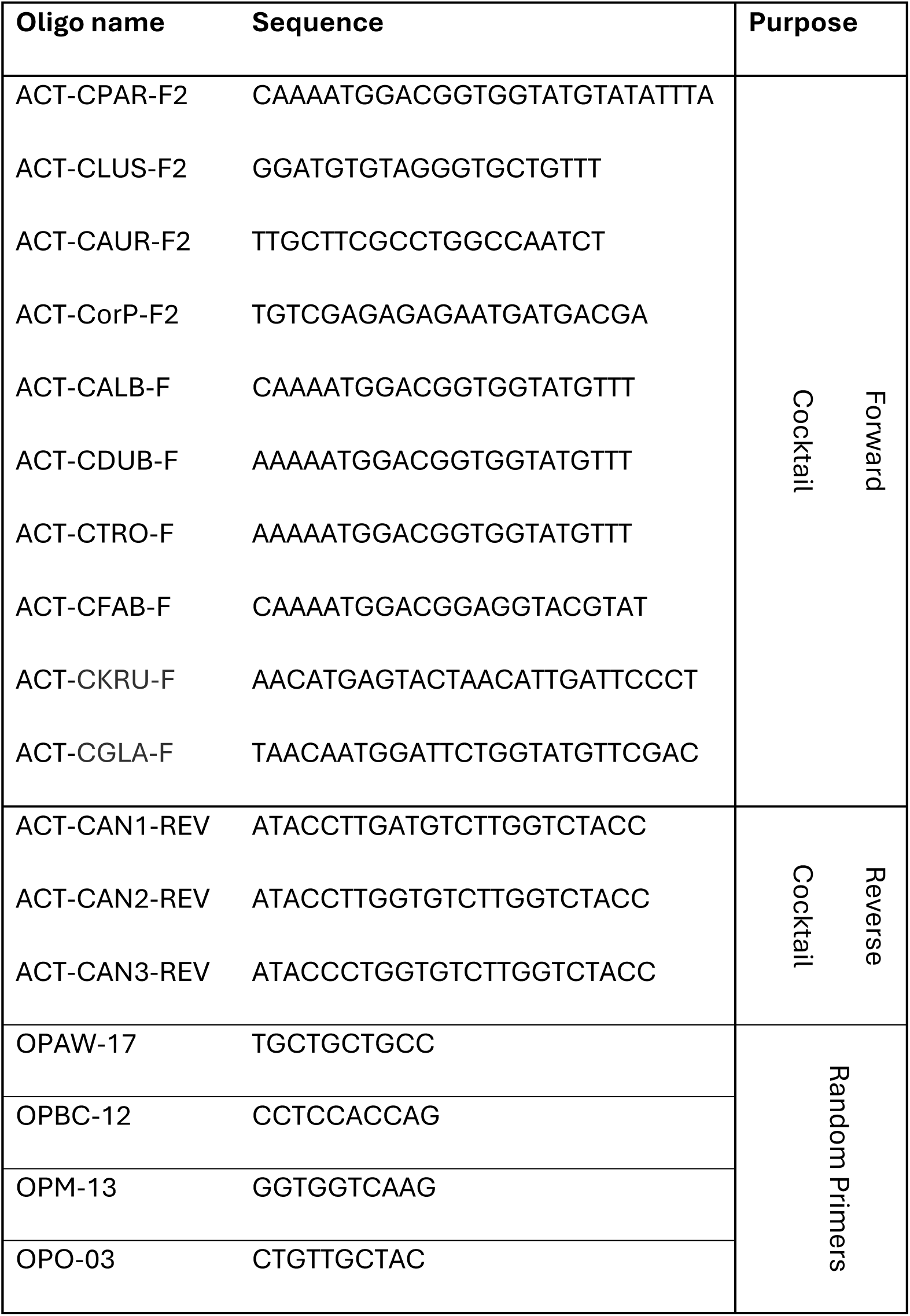
Sequence of primers used for the RAPD & CanID-PCR.

### Primer design around *ACT1* gene intron

We identified that *ACT1* gene has an intron that varies in size between species of *Candida*. We designed three reverse primers downstream of the intron, in a very conserved region among all species. Also, we designed 10 species-specific forward primers in the variable region upstream of the intron, so that the PCR sizes can differentiate between the species by approximately 50 bp. The forward and reverse primers were each pooled in an equimolar cocktail at 5 µM and 7 µM, respectively for each primer. TABLE 2 provides all details for the primers.

### PCR of *ACT1* intron for *Candida* species identification

PCR was performed using 5PRIME Hotmaster Taq DNA polymerase (QuantaBio, Beverly, MA, U.S.A.), and set up in a 25 µl reaction volume as follows: 0.5-1 µl gDNA, 1 µl each of the forward and reverse primer cocktails, 0.5 µl of 10 mM dNTP mix, 2.5 µl of 10x Taq buffer, 0.5 µl Enzyme (5U/µl), and 25 µl nuclease-free molecular biology grade water qsp. The PCR was run on a MJ Research Diad thermocycler (Biorad, Hercules, CA, U.S.A.) using the following program: (i) an initial denaturation at 94°C for 1 min; (ii) 9 cycles of : 94°C for 15 sec, 56°C for 30 sec, 68°C for 40 sec; (iii) 32 cycles of: 94°C for 10 sec, 56°C for 15 sec, 68°C for 35 sec; and (vi) a final extension at 68°C for 10 min. Alternatively PCR was performed using PFU Ultra II Fusion HS DNA polymerase (Agilent Technologies, Cedar Creek, TX, U.S.A.) or CesiumTaq Polymerase (DNA Polymerase Technology, Saint Louis, MO, U.S.A.) following manufacturer recommendations. PCR products were purified using Wizard SV Gel and PCR clean-up System (Promega, Madison, WI, U.S.A.). Purified or crude PCR products were run on a 1.5-2% agarose gel in 1x TBE for 3-6 hours.

### Alternative protocol using kits for enrichment, gDNA extraction, and PCR amplification

In the later part of the experiment, we used MolYsis™ Basic5 kit (Molzym, Bremen, Germany) which enriches for microbes by lysing host cells and digesting host DNA. gDNA from the enriched samples was then extracted using ZymoBIOMICS DNA Miniprep Kit (Zymo Research, Irvine, California, U.S.A.). We then performed PCR amplification using repliQa HiFi ToughMix (Quantabio, Beverly, MA, U.S.A.) using following program: (i) an initial denaturation at 98°C for 30 sec; (ii) 8 cycles of : 98°C for 10 sec, 55°C for 20 sec, 68°C for 15 sec; (iii) 32 cycles of: 98°C for 10 sec, 56°C for 15 sec, 68°C for 10 sec; and (vi) a final extension at 68°C for 30 sec. This alternative protocol performed well even for some samples that were not successful with the manual protocol above. Also, the repliQa PCR mix presents the advantage of being tolerant to a wide range of PCR inhibitors (Quantabio).

## RESULTS

### Primer design and testing of RAPD analysis on *Candida* colonies recovered from the positive blood culture bottles

Initially we attempted to detect polyclonal (mixed strains from same species) Candida positive blood bottles. RAPD PCR was previously shown to be able to distinguish between Candida strains within same species (8–13). So, we designed a dozen RAPD primers, among which best 4 were selected after some testing and optimization. To validate these 4 RAPD primers, we selected positive blood cultures from 4 unique patients that were identified by the clinical microbiology lab as growing single species of *C. parapsilosis*, *C. glabrata*, or *C. albicans* as identified using MALDI-TOF. We sub-cultured individual positive blood culture broth onto Sabouraud dextrose agar (SDA) plates, and selected 96 colonies from each samples to be DNA fingerprinted using RAPD. In 3 samples, the fingerprinting profiles were undistinguishable (data not shown). In the fourth sample, we found two distinct RAPD profiles on the agarose gel (Fig. 1) even though the colonies were undistinguishable on CHRCA and SDA plates. We performed Sanger sequencing of the amplified *ITS3* region of these 2 genetically different colonies and identified 2 distinct *Candida* species: *C. fabianii* and *C. parapsilosis*. Thus, using RAPD PCR, we depicted a polymicrobial (mixed species) *Candida* bloodstream infection due to identical morphotype that was missed by the clinical microbiology laboratory. Given this finding, we sought to design a quick PCR assay to identify the 10 common *Candida* species: *C. albicans, C. glabrata, C. parapsilosis, C. orthopsilosis/metapsilosis, C. tropicalis, C. dubliniensis, C. auris, C. lusitaniae* and *C. krusei*. We also included the *C. fabianii* clinical isolate. We selected the *Candida* house-keeping gene *ACT1* because of its varying size intron and a very conserved stretch just downstream of it. This allows design of PCR primers that yield PCR bands with distinct sizes for each of the species, except *C. orthopsilosis/metapsilosis* for which the PCR yields the same band size. PCR band size for each of the ten species is summarized in Table 1. To construct *Candida* species reference DNA markers, we first amplified the *ACT1* region from the 10 *Candida* reference isolates. We purified the corresponding PCR products and combined them into two size markers: **CAN Species DNA Marker I** (CAN I: *C. krusei, C. fabianii, C. albicans, C. parapsilosis* and *C. auris*) and **CAN Species DNA Marker II** (CAN II: *C. glabrata, C. tropicalis, C. dublinensis, C. lusitaniae,* and *C. meta/orthopsilosis*). By migrating the PCR product from the test sample along with *Candida* DNA markers, CAN I and CAN II, the identification of the *Candida* species in the test sample could be made based on the matching marker band size (Fig. 2).

**FIG 1.**
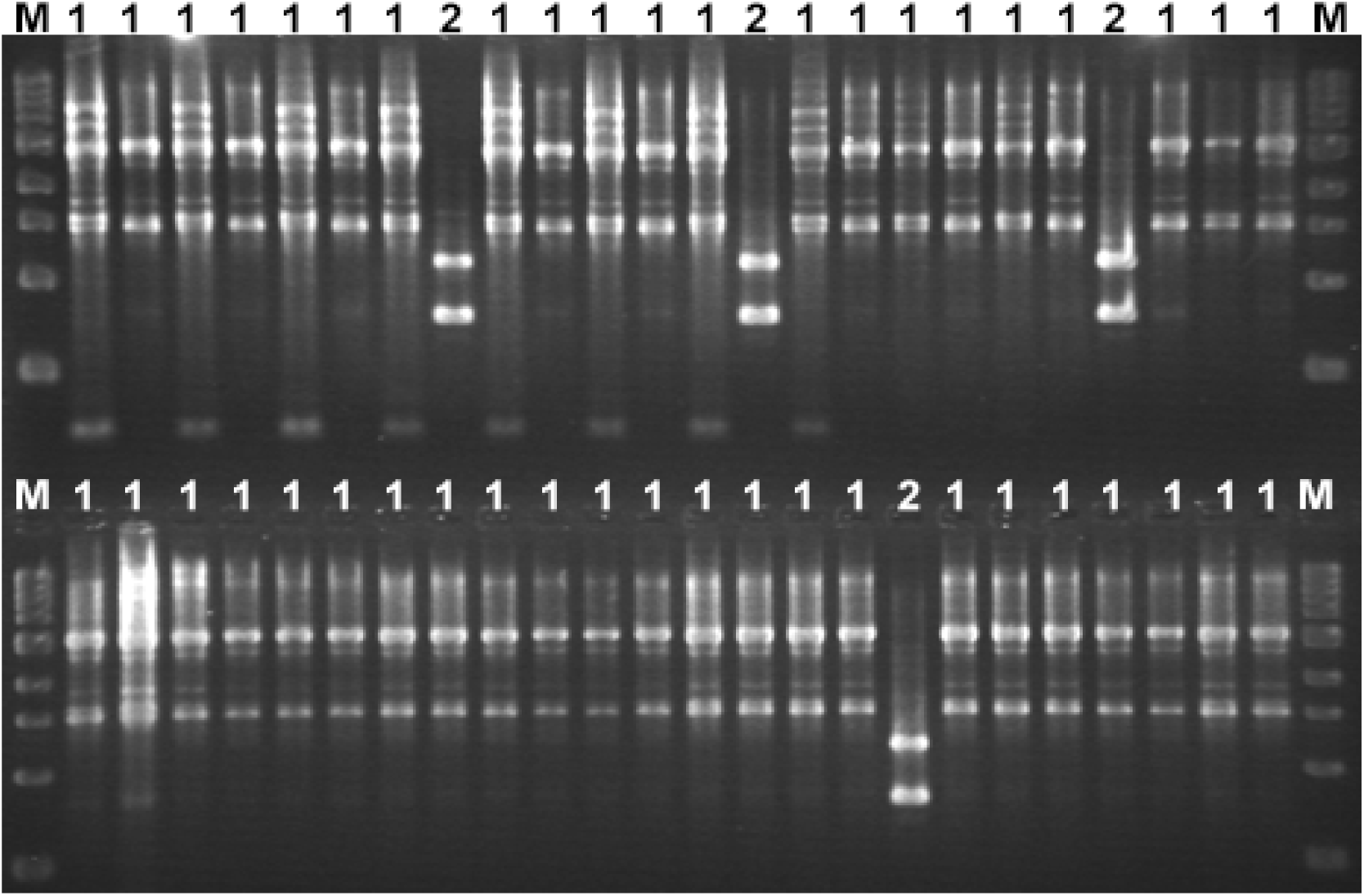
Agarose gel electrophoresis of RAPD PCR products. These RAPD profiles were obtained using primer OPAW-17 on gDNA extracted from clones (usually 96 clones, only 48 shown) isolated from Candida positive blood bottle (This gel corresponds to sample S1 shown on Fig. 3). The gel shows two RAPD profiles 1 and 2 (shown on top of each lane), which will be further identified as *C. parapsilosis* and *C. fabianii*, respectively. M is NEB 1kb DNA ladder.

**FIG 2.**
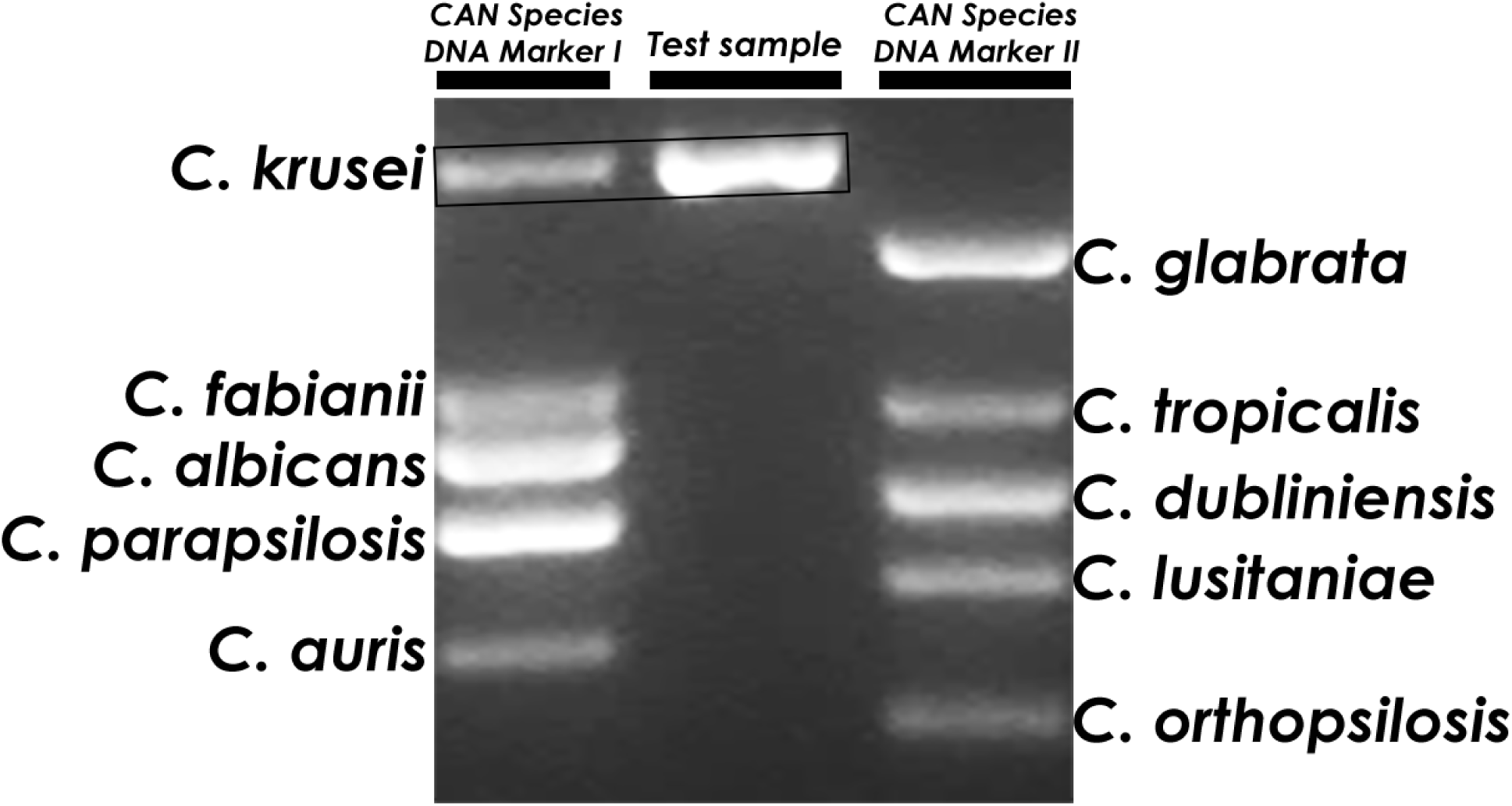
Agarose gel electrophoresis showing CanID-PCR principle. The sample’s PCR product is migrated between two lanes having each a mixture of band size markers for 5 Candida reference species, i.e. Candida species DNA Marker I or II; CAN I or CAN II, respectively (see text for more details). The sample will match one or more band size, for instance the shown sample matched *C. krusei*.

### PCR amplification and *Candida* species identification

Next, we used our assay to identify the species of *Candida* directly from the positive blood culture samples. We tested positive blood culture samples from 64 unique patients, 4 of which were used in testing RAPD on individual *Candida* colonies. Fifty-eight positive blood cultures were caused by single *Candida* spp. The most common species were *C. glabrata*, *C. albicans* and *C. krusei*, followed by *C. krusei, C. tropicalis, C. parapsilosis* complex, and one each of *C. lusitaniae* and *C. auris* (Table 3). The 6 remaining positive blood cultures were caused by 2 *Candida* species, with *C. glabrata* common in all 6; the companion *Candida* species were *C. albicans* (2 blood culture bottles), *C. krusei* (n=2) and *C. tropicalis* (n=2) (Table 3). Prior to DNA extraction, as part of our internal control, we sub-cultured the blood culture broth onto CHRCA and SDA plates. The species recovered from our subcultures matched the *Candida* species identified by the clinical laboratory in 63/64 bottles. In one bottle with dual *C. glabrata* and *C. tropicalis* identified by the clinical laboratory, CHRCA identified only *C. glabrata* while *C. tropicalis* could not be isolated.

**TABLE 3.** List of *Candida* positive blood bottle samples processed.

| **Candida species identification methods used**<br>**The number of samples identified are shown between parentheses**<br>**Percentages match with the Clinical Microbiology Laboratory are shown** |  |  |
| --- | --- | --- |
| **Clinical microbiology laboratory** | **CHROMagar Candida \*** | **CanID-PCR** |
| *C. glabrata* (n=24) | 100% (24) | 100% (24) |
| *C. albicans* (n=19) | 100% (19) | 100% (19) |
| *C. krusei* (n=6) | 100% (6) | 100% (6) |
| *C. tropicalis* (n=3) | 100% (3) | 100% (3) |
| *C. parapsilosis* (n=3) | 100% (3) | 100% (3) § |
| *C. metapsilosis* (n=1) | 0% (0) | 0% (0) |
| *C. auris* (n=1) | 100% (1) | 100% (1) |
| *C. lusitaniae* (n=1) | 100% (1) | 100% (1) |
| *C. glabrata* and *C. albicans* (n=2) | 100% (2) | 100% (2) |
| *C. glabrata* and *C. krusei* (n=2) | 100% (2) | 100% (2) |
| *C. glabrata* and *C. tropicalis* (n=2) | 50% (1) ¶ | 50% (1) ¶ |

\* Please note that CHROMagar Candida provides a presumptive identification. The result was considered a match when it agrees with the previously known speciation by MALDI-TOF MS. § One sample grew *C. fabianii* as well. ¶ Only *C. glabrata* was recovered from the sample.

PCR amplification was successful on gDNA extracted directly from blood BC. On gel electrophoresis, a single (59 samples) or double PCR bands (5 samples) of *ACT1* amplification were observed. The performance of our assay was compared with the speciation by the clinical microbiology laboratory on the same blood culture bottles. In 62 samples (97%), CanID-PCR identified the exact same species initially identified by the microbiology lab (Fig. 3 A-I); for 1 of these samples, the microbiology lab identified *C. metapsilosis*, while our tool cannot distinguish between the genetically very closely related *C. metapsilosis* and *C. parapsilosis*, as they yield exact PCR band size. They were only 2 samples with discrepant results, one (sample S1, Fig. 3A) was identified by CanID-PCR as having an extra species (*C. fabianii*), in addition to the one initially identified by the microbiology lab (C. *parapsilosis*). In the second sample (S28, Figure 3D) our PCR tool identified only *C. glabrata*, while the microbiology lab identified both *C. glabrata* and *C. tropicalis*. In total, there were 7 samples (11%) with 2 species identified by either the clinical lab or PCR, and among these, 1 of 7 samples had a species missed by the clinical lab and 1 of 7 samples had a species missed by PCR.

**FIG 3.**
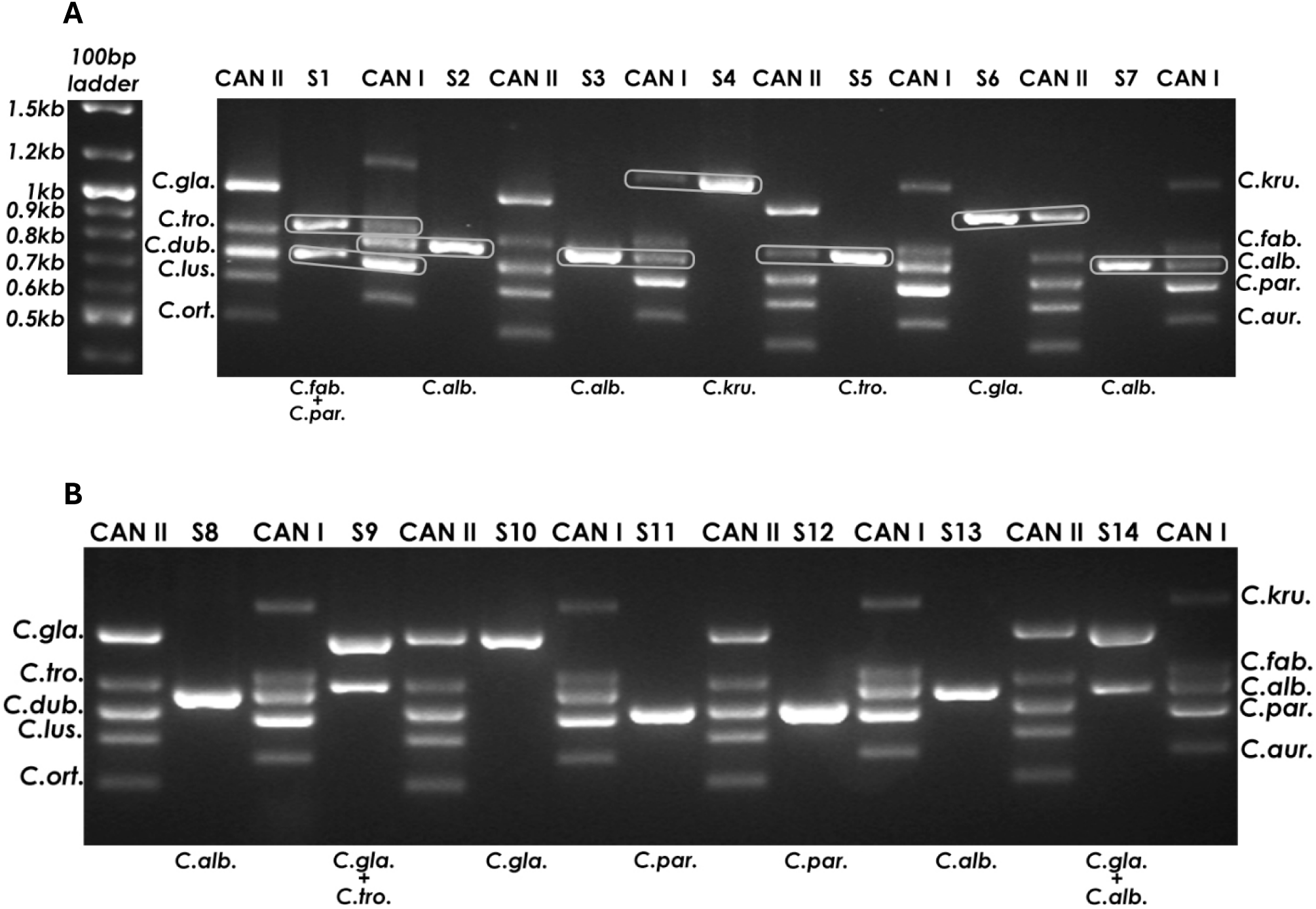
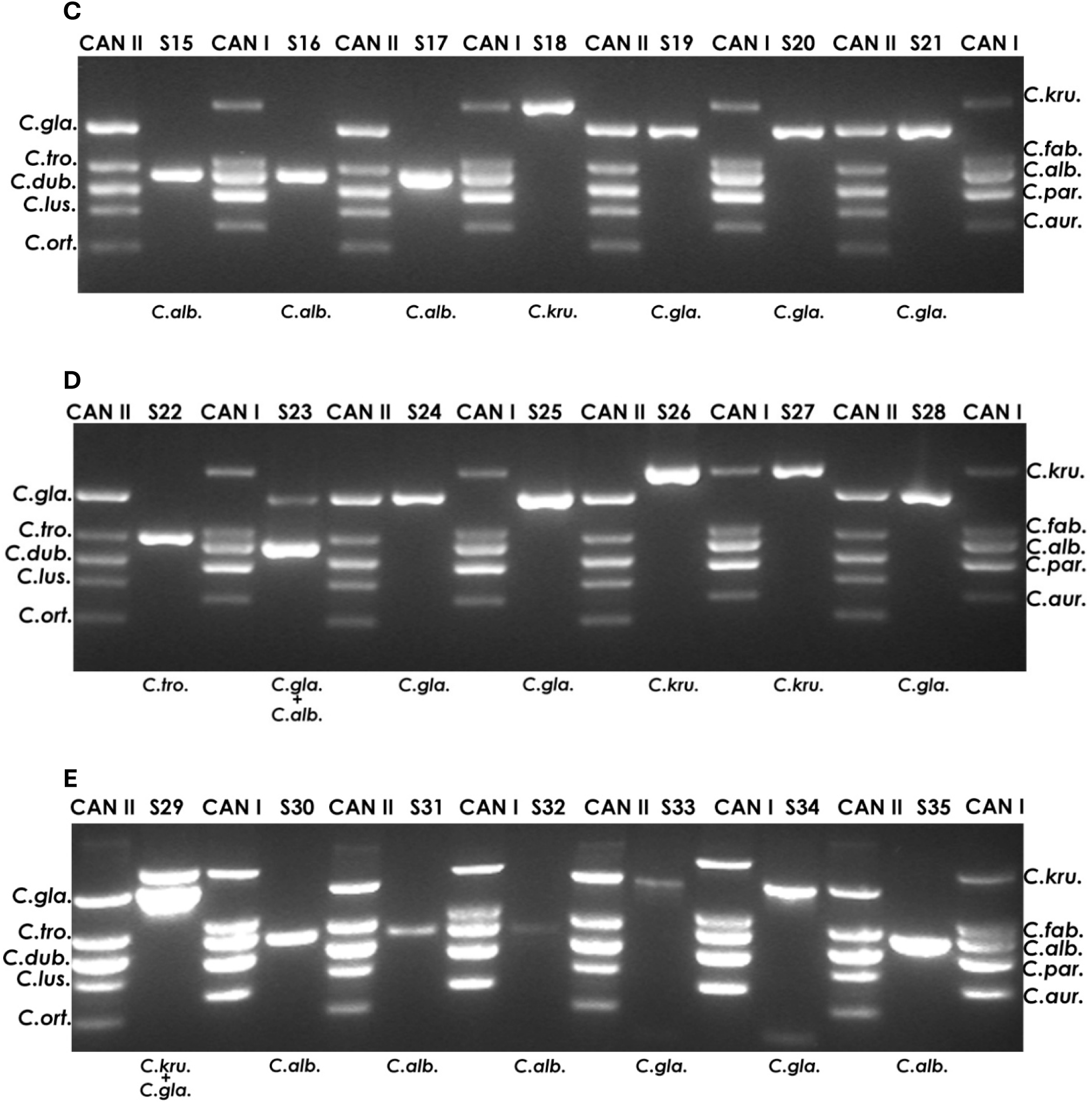
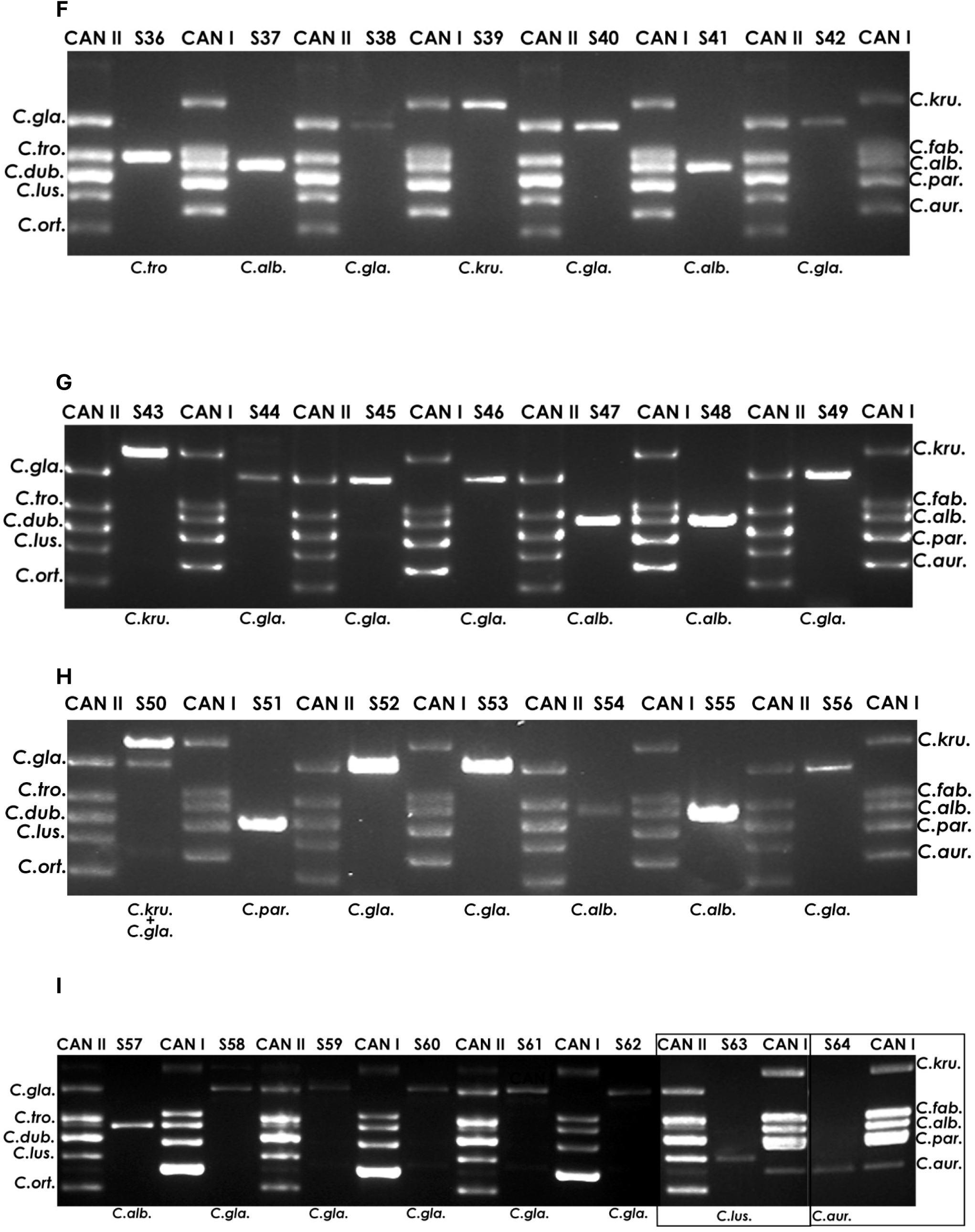
CanID-PCR results for *Candida* positive blood-bottle samples. Agarose gel electrophoresis of CanID-PCR products for 64 (panels A to I) Candida samples run along CAN I and CAN II species DNA markers. On panel A NEB 100 bp ladder is shown for reference and gray boxes indicate the species band matching for identification. All panels show a single gel, except panel I, which has 2 small gels (framed with black rectangles) attached to the main gel. The species was successfully identified on all 64 samples and indicated at the bottom of each sample’s lane. C.alb : *C. albicans*; C.kru.: *C. Krusei*; C.fab.: *C. fabianii*; C.dub.: *C. dublinensis*; C.aur.: *C. auris*; C.gla.: *C. glabrata*; C.tro.: *C. tropicalis*; C.par.: *C. parapsilosis*; C.lus.: *C. lusitaniae*; C.ort.: *C. ortho*psilosis or *C. metapsilosis*.

### Investigating discrepancies of blood culture sample S28

We investigated further the sample S28 in which the microbiology lab identified an additional species. First, we repeated the PCR and found the same result with a single band matching *C. glabrata*. We also inoculated the blood culture sample on CHRCA plates. The sample grew only *C. glabrata* in accordance with our CanID-PCR results (Fig. 4). Altogether, CanID-PCR analysis matched the species identified by the microbiology lab in 97% (62/64) of blood culture bottles.

**FIG 4.**
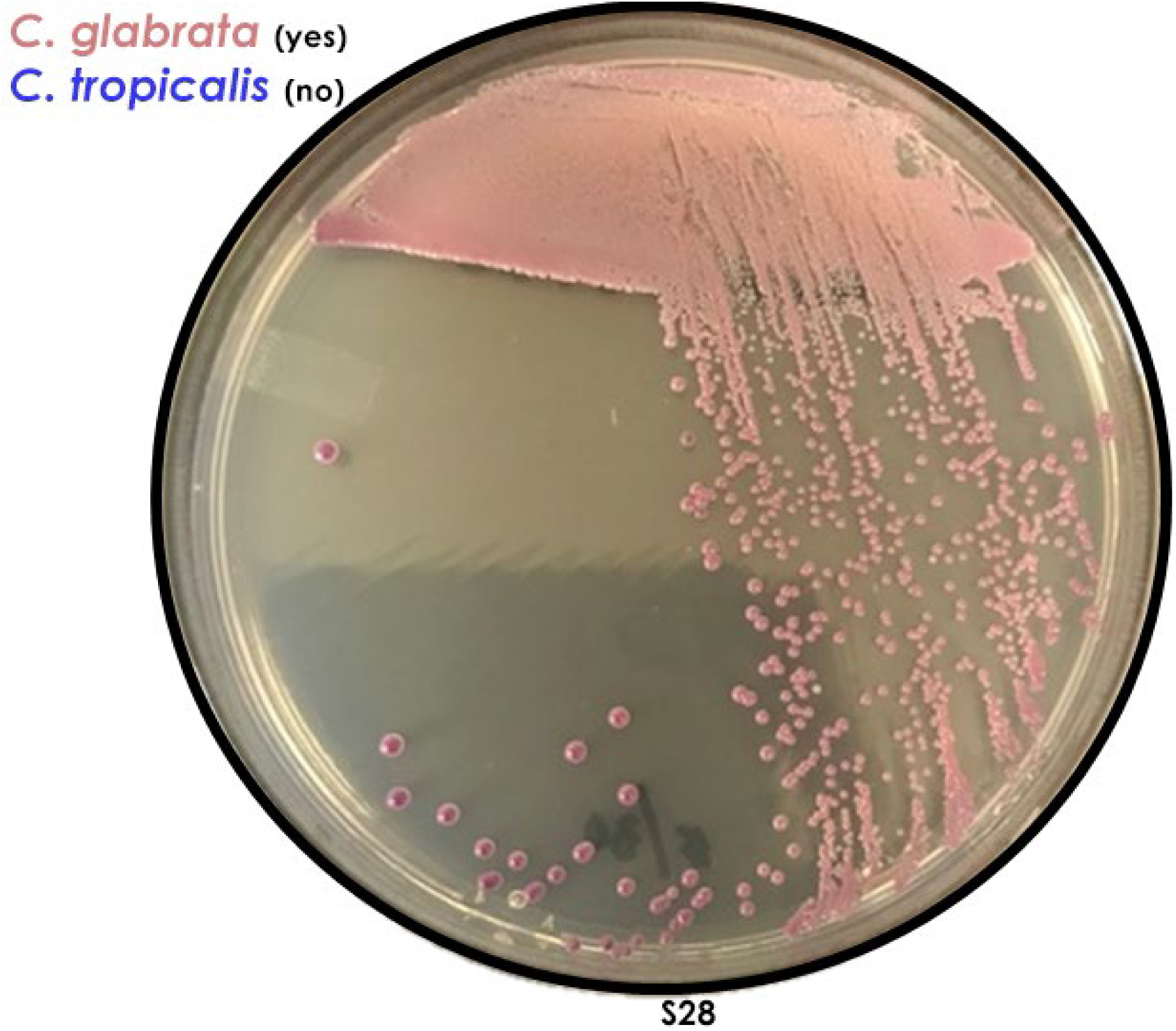
CHROMagar^TM^ Candida growth plate of sample S28. An aliquot from blood-bottle of sample S28 was re-streaked on CHRCA plate and grown for 48 hours at 30°C. Only *C. glabrata* could be identified, *C. tropicalis*, which produces distinctive metallic blue colored colonies, is not detected. This result concords with CanID-PCR.

## DISCUSSION

Our study offers 2 major findings. First, CanID-PCR analysis direct from positive blood culture bottles was able to detect with high accuracy 10 common *Candida* species causing human infections. We showed that it is a simple, quick and inexpensive technique, which can be conducted using basic laboratory equipment in as little as one day from receipt of sample to results. This technique was applied on gDNA extracted directly from blood samples which is known to present difficulty for PCR because of the presence of inhibitors (14, 15). Nevertheless, we further showed that gDNA extracted using the manual protocols or the kits were successfully amplified in all samples, although the later approach was less time-consuming and more consistent. Overall, CanID-PCR analysis accurately identified the *Candida* species identified by the clinical microbiology lab using MALDI-TOF MS in 62 of 64 samples. It is not optimized yet to distinguish between *C. metapsilosis* from *C. parapsilosis* because of their very close genetic relatedness (16). Indeed, *C. metapsilosis* belongs to the *C. parapsilosis* complex, and is phenotypically undistinguishable from other members of this complex and is less common and less virulent than *C. parapsilosis* (17). In the second sample, CanID-PCR only identified *C. glabrata* in a patient co-infected with *C. tropicalis*. We attempted to isolate *C. tropicalis* from the blood culture bottle without success and repeat PCR was not able to detect it. In further investigation, it appeared that *C. tropicalis* was identified by the microbiology lab only from the aerobic bottle and not the anaerobic bottle from which we tested with CanID-PCR. It is very likely that unlike *C. glabrata*, *C. tropicalis* was not able to grow well in the anaerobic bottle.

Our second important finding is that 9% of our patients with candidemia were infected with 2 different species. Polymicrobial candidemia has been reported to occur in ∼5% of patients, but this rate might be under-estimated because of the current workflow of the clinical microbiology lab, where only morphologically different clones were selected for susceptibility testing. In this regard, our molecular tool was able to detect a second species, *C. fabianii*, from the blood culture bottle. The clinical significance of poly-Candida species candidemia is not clear. In our case, our patient was treated with caspofungin to which both species were luckily susceptible.

In conclusion, CanID-PCR analysis directly from the positive blood cultures provides accurate speciation of *Candida* in blood, including polymicrobial candidemia. It provides speciation at least 24 hours earlier than MALDI-TOF. It is easier to perform and can be adopted easily in clinical microbiology lab for patient care as well as for epidemiologic and/or bench research studies.

